# Recursive mutational robustness in cancer through intra- and inter-genic compensation

**DOI:** 10.64898/2026.05.26.727768

**Authors:** Rohan Dandage, Aldo Hernandez-Corchado, Ariel Madrigal, Arsham Mikaeili Namini, Jichen Wang, Benedict Choi, Hani Goodarzi, Hamed S. Najafabadi

## Abstract

Mutational robustness is a fundamental property of biological systems allowing phenotypic stability against genotypic perturbation. At the gene level, this is primarily attributed to inter-genic functional redundancy driven by paralogs. Here, we show that intra-genic functional redundancy, driven by alternatively spliced isoforms, is another prevalent source of mutational robustness. Integrating isoform-resolved pan-cancer genomics and transcriptomics data across tumors and cancer cell lines, we show that alternative isoforms often bypass deleterious somatic mutations. Mutation-skipping isoforms exhibit compensatory upregulation indicative of robustness against deleterious effects, especially in highly expressed genes. We found that intra-genic robustness is strongly associated with perturbation tolerance, is highly context-specific and, notably, can be complemented and augmented recursively by inter-genic robustness. Non-tumour-suppressor genes generally show higher robustness than tumour suppressor genes, and intra-genic compensation of their mutations is associated with worse disease outcome in most cancer types. Furthermore, intra-genic compensation by mutation-skipping isoforms associates with differential isoform usage among proximal interactors, suggesting restorative protein interaction rewiring at isoform-level. Mechanistically, self-transcriptional adaptation (self-TA) following nonsense-mediated decay of perturbed isoforms provides an explanation for the observed up-regulation of mutation-skipping isoforms. Overall, the proposed recursive mutational robustness framework opens new avenues for deeper mechanistic investigation into cancer evolution and identification of selective vulnerabilities in cancer cells.

**Graphical abstract:** 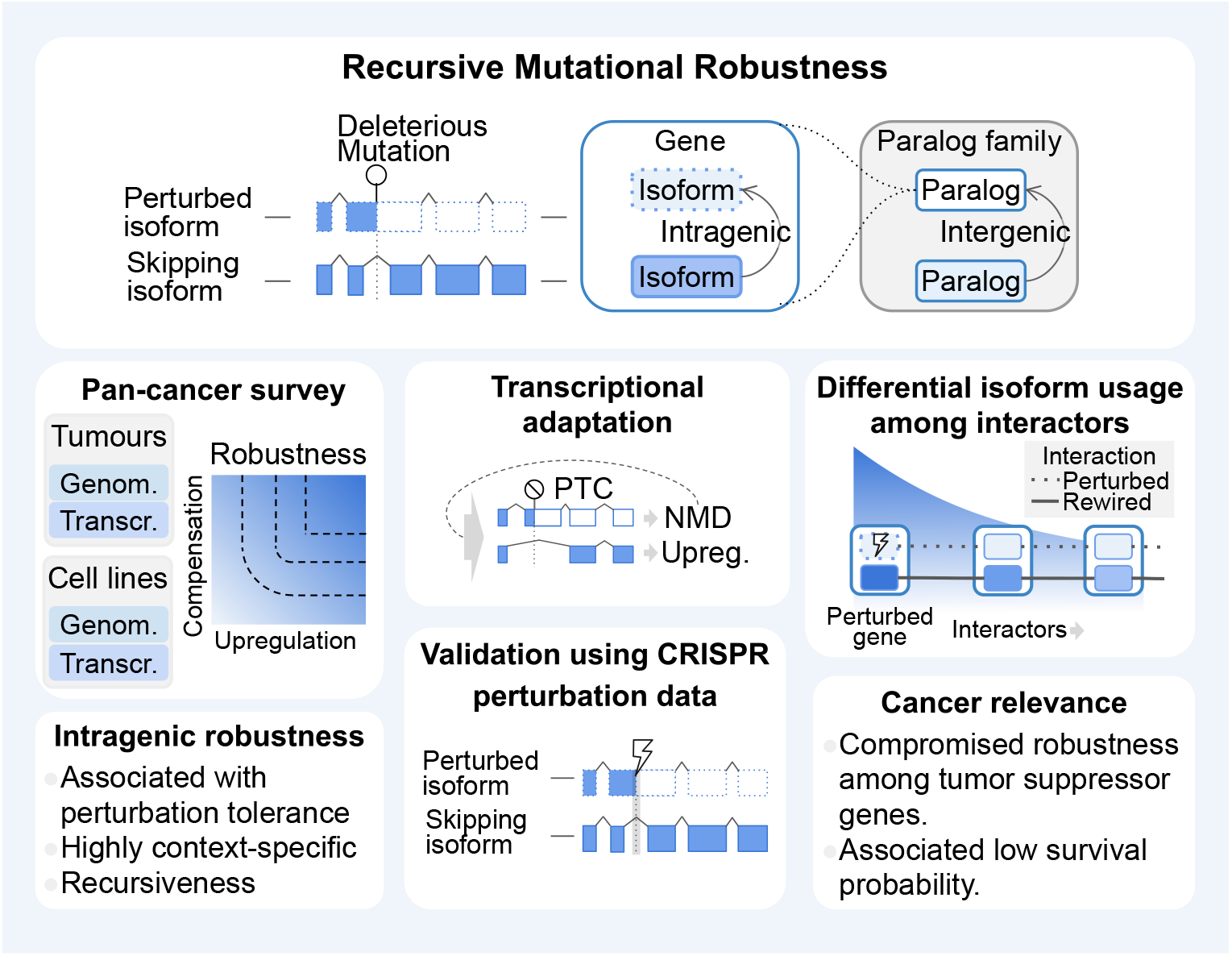

## Introduction

Cancer cells survive despite widespread genomic instability and mutational load. Previous reports ^1^suggest that nearly all coding mutations escape negative selection and are tolerated by cancer cells. This is surprising considering that many mutated genes are involved in core cellular pathways and processes. Indeed, exceptional mutational tolerance has emerged as a universal characteristic of cancer evolution ^1–4^. A partial explanation is provided by copy number amplifications that can buffer deleterious mutations ^5^. For paralogs, expression-profiling and CRISPR-based gene-deletion screens in hundreds of cancer cell lines ^6^ have revealed compensatory expression dynamics and minimal fitness defect upon deletion ^7,8^, suggesting that inter-genic robustness, via functional compensation by paralogs, is a key source of mutational robustness. However, this explanation applies mainly to highly similar paralogs, excluding genes with distant or no identifiable paralogs, which constitute a large fraction of human genes. Thus, for a large number of genes, mechanisms behind their mutational robustness remain unclear.

An unexplored potential source of mutational robustness is alternative splicing. Almost all protein-coding genes are alternatively spliced ^9^, producing isoforms that subtly diversify the gene’s function ^10,11^ and lead to protein products with nuanced yet similar molecular functions ^12,13^. Owing to this characteristic, alternative splicing is known to have consequential effects on cellular phenotypes ^14,15^ and cell differentiation ^16^, cancer initiation, progression ^17,18^, and drug resistance^19,20^. Alongside the diversification of gene function, alternative splicing also introduces functional redundancy within a gene. This redundancy is generated by the similarity of sequences, structures, and functions of the alternative isoforms of a given gene. In this respect, isoforms resemble paralogs, and indeed are dubbed as ‘internal paralogs’ ^21^, highlighting the self-similarity and recursive organisation of functional redundancy. Several studies have described the interplay between isoforms and paralogs as “function-sharing” ^22–24^.

This raises the possibility that, akin to paralogs ^25^, isoforms may serve as backup copies for each other, providing robustness against mutational perturbations. Mechanistically, this may involve expression dynamics between isoforms, resembling the dosage balance between paralogs, where the summed expression is kept roughly at a constant level ^26^ through a tradeoff between induction and attenuation ^27^. More generally, cells are known to maintain stable biomolecule concentrations through expression scaling ^28,29^ combined with tight regulations on the cell size ^30^. However, whether such dosage balance mechanism applies to isoform compensation remains unexplored. We hypothesize that isoform-level intra-genic robustness may act as a key contributor to the mutational robustness of the cancer cells. Specifically, mutation-skipping isoforms—in which perturbed regions are spliced out in matured form—could compensate for perturbed isoforms to rescue the deleterious effects of mutations. Furthermore, due to “function-sharing”, intra-genic mutational robustness may combine with inter-genic robustness, broadening the scope of influence.

Here, through a pan-cancer survey, we show that alternatively spliced isoforms not only can skip deleterious mutations, but exhibit compensatory upregulation indicative of mutational robustness. We find that intra-genic compensation of deleterious mutations associates with perturbation tolerance, is highly context-specific, and combines recursively with inter-genic compensation to confer mutational robustness. We show that transcriptional adaptation triggered by nonsense-mediated decay (NMD) explains intra-genic mutational robustness, and find that mutational robustness leads to downstream protein-protein interaction (PPI) rewiring and affects cancer outcome. Collectively, our results provide new insights into mechanisms of mutational tolerance in cancer cells.

## Results

### Pan-cancer survey of mutation skipping isoforms

Deleterious mutations, such as a stop gain in a coding exon, can render an isoform dysfunctional. However, there is a possibility that alternatively spliced isoforms can skip the mutated region in the mature form (**Fig 1A**). To assess the frequency of this phenomenon and whether the mutation-skipping isoforms tend to be expressed, we carried out a survey across 30 tumor types ^31^ and cancer cell lines representing 21 distinct lineages ^32^ using genomics and transcriptomics data (**Fig 1B**). Using the whole genome sequencing data, we first sought to identify isoforms carrying deleterious mutations that are highly likely to cause functional perturbation. For this purpose, we developed a classification system that marks mutationally perturbed isoforms (see **Supplementary Methods**), overcoming the limitation of commonly used gene-centric methods. We first carried out a pan-cancer survey of possible functional relevance of the mutation-skipping isoforms. We found that the fraction of a gene’s isoforms that skip deleterious mutations and get expressed is comparable to the fraction of isoforms a gene typically expresses (median percentages 33.33 vs 39.22, **Fig 1C**-left), indicating that the expression of skipping isoforms is common. Across all tumour types as well as in cancer cell lines, such mutation-skipping and expressed isoforms are more likely to undergo exon-skipping (SE, **Fig 1C**-middle, two-sided Mann-Whitney U-test, FDR<0.05 Benjamini-Hochberg), suggesting that cassette exon skipping is the predominant form of alternative splicing that creates mutation-skipping isoforms. Overall, this survey revealed that mutation skipping by isoforms is common, mutation-skipping isoforms tend to be expressed, and that exon-skipping is the primary alternative splicing event leading to mutation-skipping.

**Figure 1:**
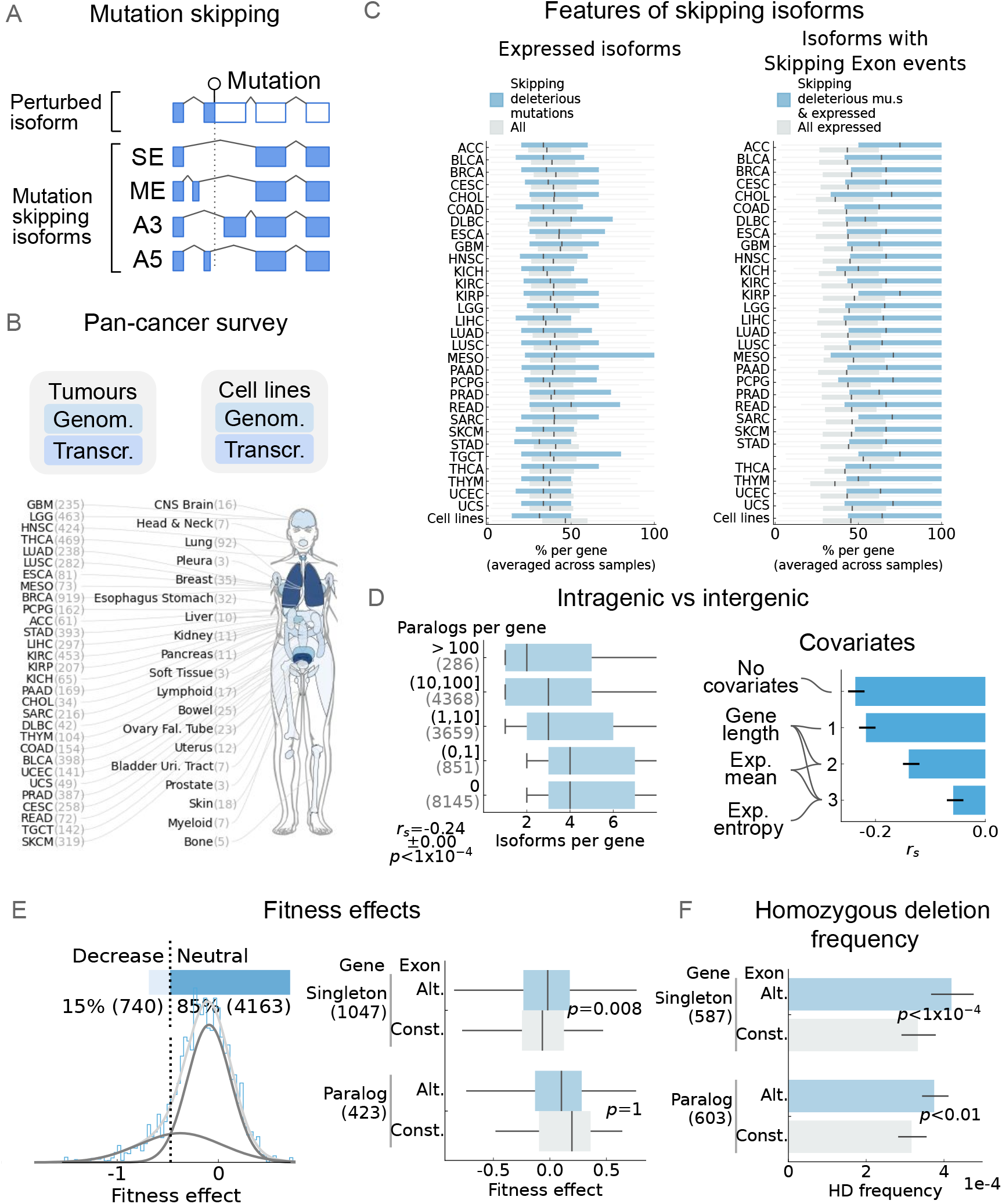
Pan-cancer survey of mutation skipping isoforms and its functional consequences. **A**: Schematic presentation of the mutation-carrying and -skipping isoforms. Mutation skipping can occur by skipping a cassette exon (SE), usage of mutually exclusive exon (ME) or skipping a part of the exon (A5: alternative 5’ or A3: alternative 3’) where the mutation is located. Although not depicted, depending on the location of the mutation, mutation-skipping can also occur with other splicing events such as AF (alternative first) and AL (alternative last). **B**: Overview of our pan-cancer survey, showing the cancer types of tumors and the lineages of the cell lines analyzed and the corresponding number of samples. **C**: Properties of the mutation-skipping isoforms across cancer types represented by tumors and cell lines. For each of the properties, percentage per gene was calculated in a sample-wise manner and then averaged over the corresponding sample subset. See **Methods** for details. Left: Percentage of isoforms skipping deleterious mutations and expressed (TPM>1) is compared against the percentage of (any) expressed isoforms per gene. Right: The percentage of isoforms with SE events among isoforms skipping deleterious mutations and expressed (TPM>1) is compared against that of any expressed isoforms per gene. **D**: Left: the number of isoforms (x-axis) per gene is correlated with the number of paralogs per gene (y-axis). *r*_*s*_: Spearman’s correlation coefficient, *p*: *p*-value associated with the correlation. Right: the effect of step-wise inclusion of covariates (y-axis) on the partial correlation coefficients between the number of isoforms and number of paralogs per gene (Spearman, x-axis) is shown. The error bars indicate CI95%. **E**: Fitness effects of mutation skipping. Left: distribution of the fitness effects of exon deletions obtained from the genome-wide CRIPSPR-based screen (data from ^35^), along with the percentages of the exons whose deletions led to decrease in fitness and those that had neutral effect (top). The distribution (histogram in blue) is fitted with a gaussian-mixture model (light gray). The intersection between the inferred sub-distributions (gray) is used as a threshold (dotted line) to classify the exons with fitness decrease and neutral effects. Right: the fitness effects are compared between alternative (Alt.) and constitutive (Const.) exons, for the singleton and paralogous genes. *p*: *p*-value obtained from one-sided Mann-Whitney U-test. In this figure (and all following boxplots), the central line indicates the median, the extent of the box is from the first quartile (Q1) to the third quartile (Q3), and the whiskers extend to Q1-1.5×IQR and Q3+1.5×IQR. The sample sizes are indicated in the parenthesis. **F**: Homozygous deletion (HD) frequencies are compared between alternative (Alt.) and constitutive (Const.) exons, for the singleton and paralogous genes. The bar lengths indicate mean value, and error bars indicate 95% confidence interval. *p*: *p*-value obtained from one-sided Mann-Whitney U-test.

Considering the similarity of the functional redundancy between isoforms and paralogs, next, we explored the interplay between inter-genic and intra-genic functional redundancy. We observed a strong anticorrelation between the number of paralogs and isoforms per gene (**Fig 1D**-left), suggesting that inter-genic and intra-genic functional redundancy may have complementary roles. Consistent with this notion, gene-level dispensability—obtained from a large-scale CRISPR based gene-deletion screen ^33^—correlated with high paralog numbers, underlining robustness to whole-gene inactivation conferred by paralogs. In contrast, gene-level essentiality correlated with high isoform numbers (**Fig S1A**), supporting the notion that inactivation of high-isoform genes is rarely if at all compensated by the functions of other homologous genes. We inspected the covariates of the anticorrelation between the numbers of isoforms and paralogs (**Fig S1B**), and identified normalized expression entropy, i.e., the variability in which isoforms are expressed, to have a strong influence (partial spearman correlation r_s_reduced to –0.06), alongside known factors such as gene length and gene-level expression level ^34^ (**Fig 1D right**).

Interestingly, while gene-level inactivation of high-isoform genes is deleterious, exon-deletion is often tolerated. From a large-scale CRISPR-based exon-deletion screen dataset (>12,000 exons in two cell lines) ^35^, we found that, in genes with pronounced gene-level fitness effects (see **Supplementary Methods**), deletion of ∼85% of exons had a neutral effect (classified using Gaussian mixture modeling of the distribution of fitness effects; **Fig 1E**-left), suggesting that dispensability is common. Among singleton genes, i.e., those without identifiable paralogs, deletion of constitutive exons has significantly larger fitness effects compared to alternative exons (*p*=0.008), whereas in genes with paralogs this difference disappears (*p*=0.98; **Fig 1E**-right). These observations suggest that, while genes without paralogs are often essential, perturbation of their alternatively spliced exons is relatively tolerated, underlining the possibility of isoform-driven robustness.

We orthogonally examined this trend by using homozygous deletion (HD) frequency between the exon types. The HD frequency reliably measures perturbation tolerance because of the complete loss of genomic region, as shown before ^36^. Using HD frequencies from the tumor dataset, we found that the alternatively spliced exons tend to have a greater tolerance than constitutive ones (**Fig 1F**). This trend, unlike the CRISPR-based exon fitness analysis, was also seen among paralogs (*p*=0.001), but the effect was substantially stronger in singleton genes (*p=*4e-6). Together, these observations suggest that perturbation of alternative exons is widely tolerated, suggesting the possibility that alternative isoforms can compensate for each other. Collectively, the pan-cancer expression of mutation-skipping isoforms, and the high tolerance of alternative exons to perturbation provide strong evidence for intra-genic functional redundancy. Cancer cells exploit this redundancy to tolerate deleterious mutations through functional compensation, as we explore in the next section.

### Mutation-skipping isoforms are frequently up-regulated in response to somatic mutations

To test if mutation-skipping isoforms functionally compensate for those perturbed by deleterious mutations, we leveraged the dosage balance principle, specifically the notion of “backup upregulation”, which has been well established at the inter-genic level ^37^: considering a gene perturbation, where all isoforms of a gene are mutationally perturbed, inter-genic mutational robustness may manifest as the compensatory upregulation of paralogs that, in practice, bypass the mutation (**Fig 2A**-left). We examined whether similar backup circuits exist intra-genically, i.e., whether a similar compensatory upregulation of mutation-skipping isoforms can be detected in tumors or cancer cell lines harboring deleterious mutations affecting specific isoforms, using their genome and transcriptome sequencing data. To make the connection between inter- and intra-genic backup circuits more explicit, we first defined a common terminology where “entity” refers to isoforms intra-genically and paralog genes inter-genically, while “group” refers to genes intra-genically and paralog families inter-genically (**Fig 2A**-right). The mutational robustness scores are ascribed to the interaction between the perturbed and the skipping entities. Due to the vast combinatorial space of possible interactions, we restricted our analyses to entities with high functional similarity (See **Supplementary Methods**).

**Figure 2:**
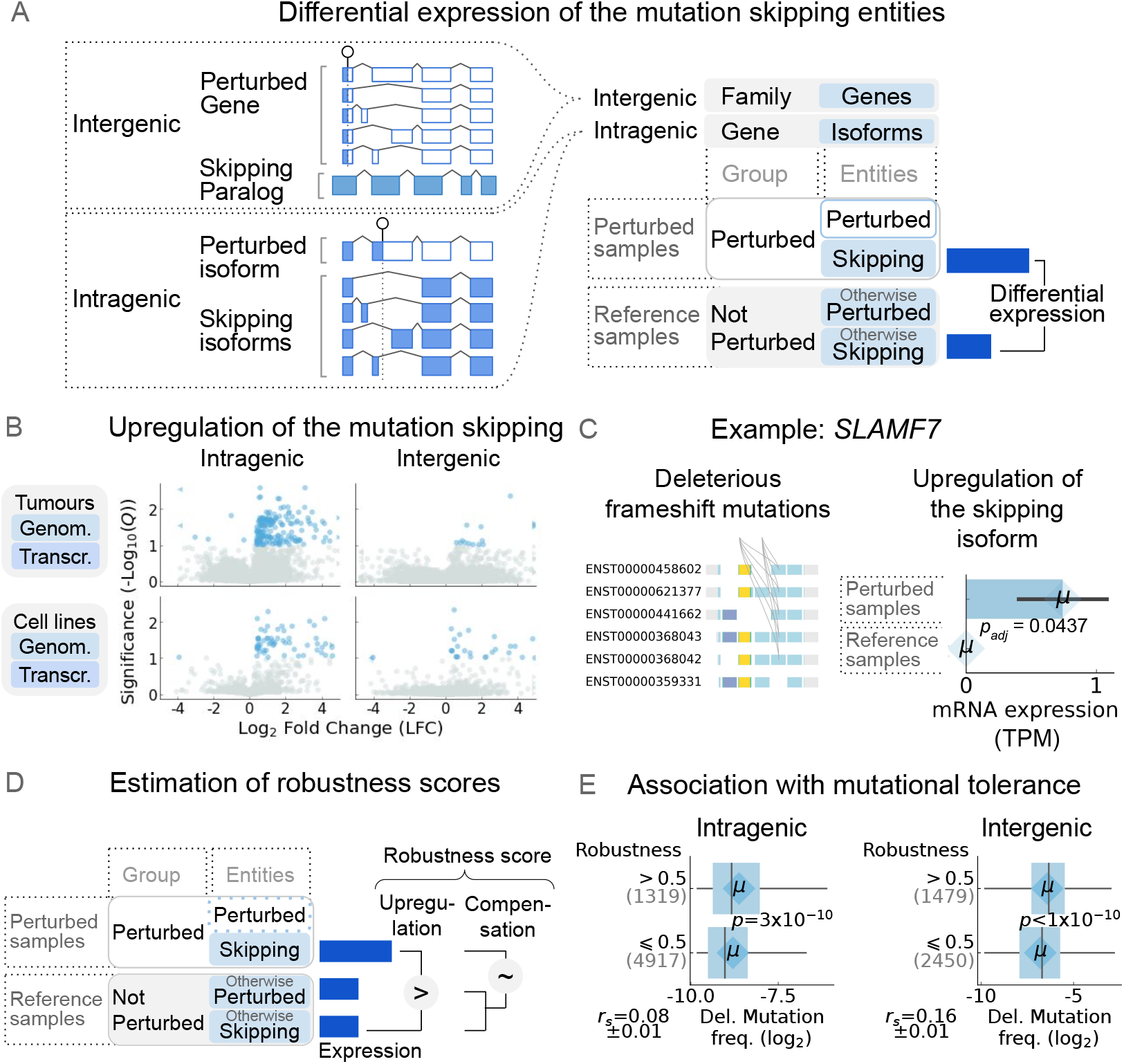
Mutation robustness at the intra-genic and inter-genic levels. **A**: Left: schematic representation showing intra- and inter-genic levels. For the inter-genic case, a perturbed gene and its paralog, which remains unperturbed, is shown. For the intra-genic case, perturbed and mutation-skipping isoforms within the same gene are shown. Right: the terminologies are indicated. **B**: Upregulation of the mutation-skipping isoforms (left) and genes (right) in the tumor (top) and cell line (bottom) datasets, shown as volcano plots. The points shown in blue indicate statistically significant hits (FDR<0.1). *Q*: Adjusted *p*-value obtained from differential expression analysis. **C**: An example gene, *SLAMF7*. For the isoform that skipps the deleterious frameshift mutations (left), abundance between samples with and without mutation are shown (right). Error bars indicate standard deviations. *p*_*adj*_: adjusted *p*-value obtained from the differential expression analysis. **D**: The mutational robustness scores were estimated by combining two measures: (i) upregulation (LFC>0 based on differential expression) and (ii) compensation (ratio of skipping entity expression in perturbed samples to total group expression in reference samples). The two scores are then integrated into a single robustness score (see **Methods** for more details). **E**: Association of the mutational robustness scores (y-axis) with deleterious mutation frequency serving as the measure of perturbation tolerance (x-axis) at intra- (left) and inter-genic (right) levels. *p*: *p*-value obtained from two-sided Mann-Whitney U-test; *r*_*s*_: mean Spearman’s correlation between continuous values.

We expectedly observed frequent upregulation of mutation-skipping (backup) entities at the inter-genic level (16 and 25 significant up-regulation events in tumors and cell line datasets, respectively, at FDR<0.1; **Fig 2B**-right). Intriguingly, we found an even stronger trend at the intra-genic level, in both the tumor and cell line datasets (162 and 59 significantly up-regulated mutation-skipping isoforms in tumors and cell lines, respectively, at FDR<0.1; **Fig 2B**-left). This stronger intra-genic effect likely stems from the generally greater functional similarity of isoforms compared to paralogs. In contrast, downregulation of the mutation-skipping isoforms is rare (only six instances in tumor and one in cell lines at FDR<0.1), suggesting an overall directional trend where mutations affecting a subset of a gene’s isoforms mostly lead to up-regulation of isoforms skipping those mutations (**Fig 2C** shows an example). Apart from restricting the combinatorial search space, the above analysis relies on statistical testing using a limited number of samples. Consistent with the well-known “long tail” of rarely mutated genes ^38^, most genes in our analysis carried deleterious mutations in one or a few samples (**Fig S2**), which were then compared to a few non-perturbed tumors or cell lines with closely matching transcriptome-wide expression profiles. Thus, each differential analysis involves comparison of a few samples within each of the perturbed with a corresponding set of non-perturbed reference samples, limiting statistical power of the analysis. Therefore, we note that the number of up-regulated mutation-skipping isoforms that we observed is likely a substantial under-estimation of the true number of cases.

We next estimated the mutational robustness as a combination of (i) the extent of backup upregulation of the mutation-skipping entities, and (ii) the extent to which the mutation-skipping entities compensate for the loss of the perturbed entity (**Fig 2D**, see **Methods** for details). Briefly, the upregulation was measured by differential expression analysis – using limma ^39^ – comparing the expression of the skipping entities between samples with the perturbation (i.e., samples with a mutation in the perturbed entity) and a set of reference samples that lacked any mutations in the perturbed entity and had, overall, a comparably similar genome-wide expression profile to that of the perturbed samples (see **Methods** for details). Then, to assess the extent to which upregulation can amount to compensation for the loss of perturbed entity, a compensation score was calculated as the ratio between the upregulated expression in the perturbed samples and the cumulative expression of the group in the reference samples. Finally, the upregulation and compensation scores were combined in a unified robustness score (see **Supplementary Methods** for details on the calculation of robustness score). To explore the extent to which backup isoform up-regulation confers mutational robustness, for entities with log fold change (LFC) >0, we calculated the compensation score followed by the calculation of a robustness score, whereby we combine the LFC and compensation score (**Fig S3A**). We used the score threshold of 0.5 to identify instances with strong robustness (**Fig S3A**), and in the subsequent analyses we pooled together these robustness classifications across tumor and cell line datasets to increase statistical power (separately for intra- and inter-genic interactions).

Several lines of evidence support the relevance of this stratification based on our calculated robustness scores. First, we observed that strong intra-genic mutation robustness was significantly associated with high similarity between perturbed and mutation-skipping isoforms (OR=1.55, p=2e-8; **Fig S3B**); this relationship was less clear for inter-genic mutational robustness (OR=1.1, *p*=0.66). Also, both intra- and inter-genic robustness are strongly associated with smaller group sizes (i.e., genes with <10 isoforms, OR= 1.21 with *p*<1e-10, and paralog groups with <10 paralogous genes, OR=1.48 with *p*=2e-9; **Fig S3B**). Together, these observations suggest that robustness tends to be more common among less diverged, smaller groups, as expected. Second, we found that strong robustness is significantly associated with independent measures of tolerance to perturbations. Specifically, deleterious mutation frequency in tumours, which represents the degree to which such mutations are tolerated, is strongly associated with robustness (p<3e-10 for both intra- and inter-genic robustness; **Fig 2E**). Tolerance of CRISPR-based perturbations showed a similar trend in terms of association with robustness (using exon perturbation data from ref ^35^ for intra-genic and gene perturbation data from ref ^40^ for inter-genic; shown in **Fig S4**), although this association did not reach statistical significance based on exon perturbation data. Together, these observations suggest that robustness scores obtained from transcriptomic analysis of tumour and cell line data mirror orthogonal measures of perturbation tolerance.

### Mutation robustness and recursiveness between intra and inter-genic levels

Our pan-cancer analysis also revealed an important contrast between the intra and inter-genic levels. Intra-genic mutational robustness instances were more context-specific (normalised entropy, *Ĥ*=0.6)—often occurring in one cancer type—compared to inter-genic ones (*Ĥ*=0.8, **Fig 3A**). We used the tumor dataset for this analysis given its broader coverage of cancer types. This trend may be driven by the highly context specific nature of isoform-level expression (**Fig S1A**-left).

**Figure 3:**
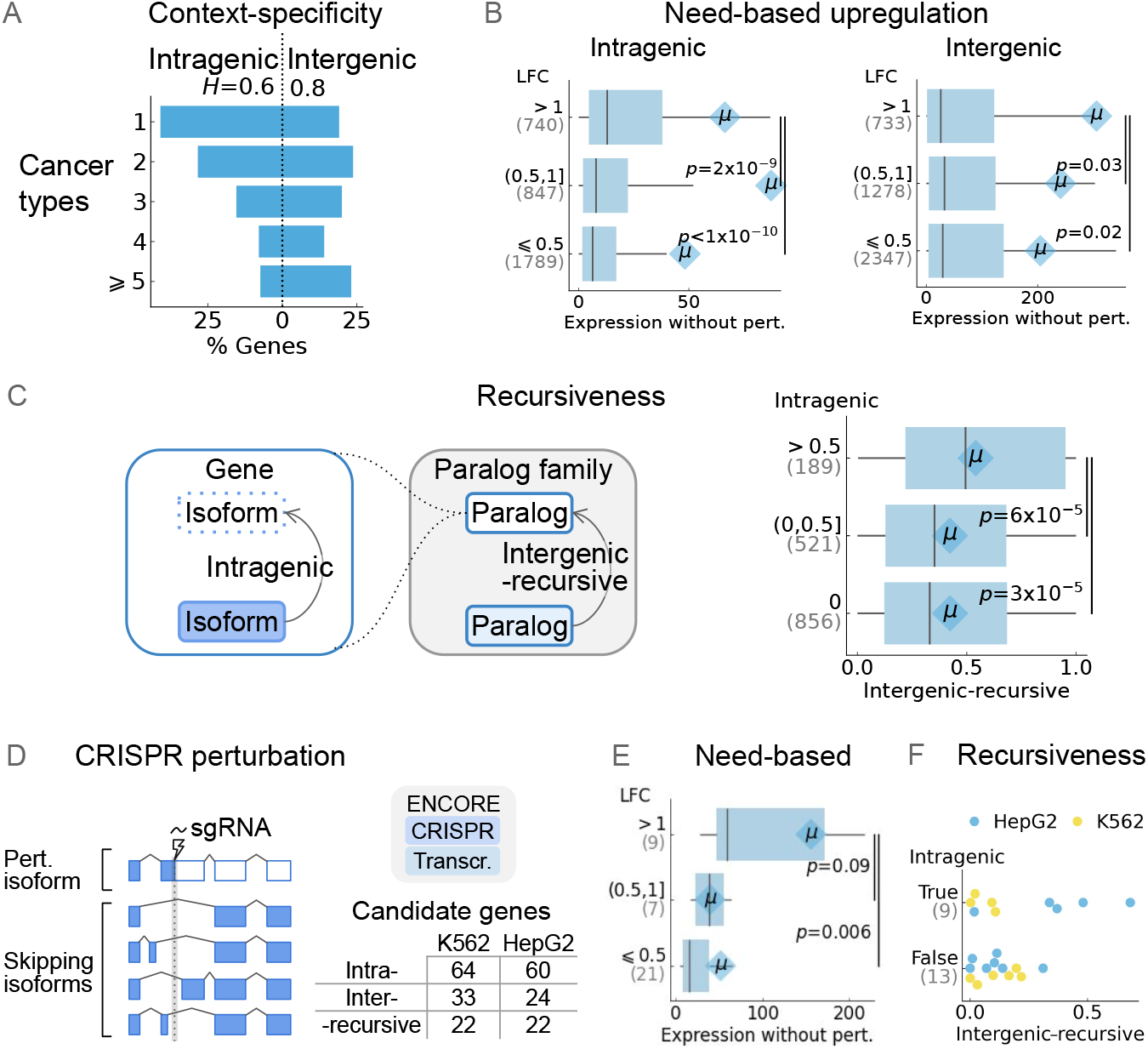
Need-based and recursive nature of mutational robustness. **A**: Context-specificity of the identified robust genes (robustness score>0) is visualised as distribution across the number of cancer types (in the tumor dataset) they were identified in. *Ĥ*: normalised entropy. **B**: Need-based upregulation tested by comparing the LFC of differential expression (y-axis) against the expression (TPM, x-axis) of isoforms (intra-genic, on top) and genes (inter-genic, bottom) in unperturbed reference samples (see **Methods**). Due to the long-tailed distributions of expression values, mean values are indicated in the diamond shapes. *p*: *p*-value obtained from two-sided Mann-Whitney U-test. **C**: Recursiveness of mutational robustness. On the left, schematic presentation of the ‘recursive’ setting. On the right, inter-genic scores are plotted across bins of intra-genic scores. *p*: *p*-value obtained from two-sided Mann-Whitney U-test. **D**: Schematic presentation of CRISPR-Cas9-perturbed isoforms and those skipping the perturbation. The sgRNA location is depicted as a dotted line; the gray area shows the average span of perturbation. Mutational robustness scores were estimated using transcriptomic profiling of CRISPR-perturbed genes from ENCODE. The number of candidate genes for estimating intra-genic robustness scores is shown. Note that those scored for recursive robustness are the intersection of intra- and inter-genic cases. **E**: Need-based upregulation shown using the plot similar to panel B, but based on CRISPR data. **F**: Inter-genic robustness scores are compared between isoform perturbations which exhibited *any* intra-genic robustness (score>0) versus those that did not (score=0).

Probing at the functional importance of mutational robustness exhibiting entities, we tested the concept of need-based upregulation which has been established in the case of inter-genic functional redundancy ^41^, meaning upregulation is likely to occur in contexts where gene function is necessary. Specifically, we examined whether robustness is higher in contexts where the baseline expression of the gene is higher. For both intra- and inter-genic mutational robustness, consistently we find this trend to hold true, with a relatively stronger association at intra-genic level (*p*<1e-10 for intra) than inter-genic level (*p*=0.02 for inter, **Fig 3B**). This indicates that the genes exhibiting robustness tend to be highly expressed, with potentially essential molecular functions ^42^. Indeed, the enrichment of important molecular functions among the intra-genic and inter-genically robust genes provides further support for the notion of need-based upregulation (**Fig S5**-left)— we observed that functions enriched at intra-genic level are involved in maintenance of regulatory networks and genomic architecture necessary for controlled growth and replication (e.g., chromatin binding, protein C-terminus binding) whereas those enriched at inter-genic level mediate critical interactions with the external environment and signal transduction pathways essential for cellular adaptation and homeostasis (e.g., G protein-coupled receptor activity, transmembrane transporter activity). The functional “need” was also evident from the enrichment of essential biological processes (e.g.,housekeeping processes such as lysosomal acidification and cell projection assemblyc) and cellular components (e.g., cytoskeleton and nucleosome, **Fig S5**-middle and -right), after accounting for the confounding effects of covariates known to associate with the genes with many isoforms and paralogs (**Fig S1A**, see **Supplementary Methods**).

We also explored the possibility that genes could simultaneously be subject to inter- and intra-genic mutational robustness. In this scenario, a mutation that affects only a subset of isoforms is compensated through up-regulation of mutation-skipping isoforms as well as up-regulation of paralogs. We categorise this setting as ‘recursive’ since the same compensation framework repeats itself in two nested groups: within the paralog group, the perturbed entity is a gene, which is compensated by other genes within that group, and at the same time, within the gene, the perturbed entity is a subset of isoforms, which is compensated by other isoforms of the same gene (**Fig 3C**-left). To assess the mutational robustness in this recursive setting, we estimated inter-genic mutational robustness for the subset of genes for which intra-genic mutational robustness could be computed. We observed that, in the presence of strong intra-genic robustness, inter-genic robustness is significantly larger (**Fig 3C**-right, *p*=3e-5), suggesting that the two mechanisms can further augment each other. This result suggests that the intra-genic level of mutational robustness is not isolated, but is often accompanied by inter-genic mutational robustness.

Interestingly, need-based and recursive robustness are not only observed for somatic mutations in cancer, but also for engineered mutations. Specifically, we systematically analyzed CRISPR-Cas9-based perturbations from ENCODE^43^ (representing mostly RNA-binding proteins but also other genes), where CRISPR perturbation was carried out using single sgRNAs in K562 and HepG2 cell lines. For many targeted genes, we found that the perturbed region – spanning sgRNA target +/-50 bp on average ^44^ – could potentially be spliced out in some of the isoforms of the gene (**Fig 3D**). Such perturbation-skipping in some isoforms made the corresponding genes candidates for the estimation of intra-genic robustness scores. We found seven genes showing significant upregulation of perturbation-skipping isoforms following CRISPR perturbation relative to control cells (FDR<0.1, **Fig S6A**). For example, in the case of the *CAST* (Calpastatin), among the isoforms which skipped the sgRNA targeting, the one with the least affected inhibitory domain showed upregulation in expression in perturbed samples (**Fig S6B**). Overall, we observed that genes that show strong intra-genic compensation response to CRISPR perturbations are expressed, at baseline, at significantly higher level compared to genes that did not show compensatory up-regulation of perturbation-skipping isoforms (*p*=0.006, **Fig 3E**), mirroring the pattern of need-based upregulation we observed in somatic mutations and. This observation further supports the notion that highly expressed, potentially functionally important genes are more likely to exhibit robustness. Also, aligning with our observation for somatic mutations (**Fig3C**-right), we found that genes exhibiting intra-genic compensation of CRISPR perturbations are also more likely to exhibit, at the same time, strong inter-genic compensation by up-regulation of paralogs (**Fig 3F**), further underlining the recursiveness of mutational robustness (specific examples are shown in **Fig S7**).

### Splicing shift and self-transcriptional adaptation explain intra-genic mutational robustness

To investigate the mechanisms of mutational robustness, we first assessed the features of the relevant deleterious mutations. We mainly focused on the intra-genic level, which remains largely unexplored. First, focusing on splicing-related features, we found that, uniquely at the intra-genic level, splice-site deleterious mutations were associated with higher robustness scores compared to non-splice-site mutations (*p*=4e-9, **Fig 4A**-left). These associations are consistent with the expectation that mutations disrupting splice sites would be more likely to result in splicing-mediated isoform switch. We also saw a similar trend in exonic mutations in close proximity of splice sites (≤2 nucleotides away from splice site), which showed significantly higher intra-genic robustness (but not inter-genic robustness) compared to more distal exonic mutations (*p*<3e-4, **Fig 4A**-middle), more so for donor site-proximal mutations compared to acceptor site-proximal mutations (*p*=0.01, **Fig 4A**-right).

**Figure 4:**
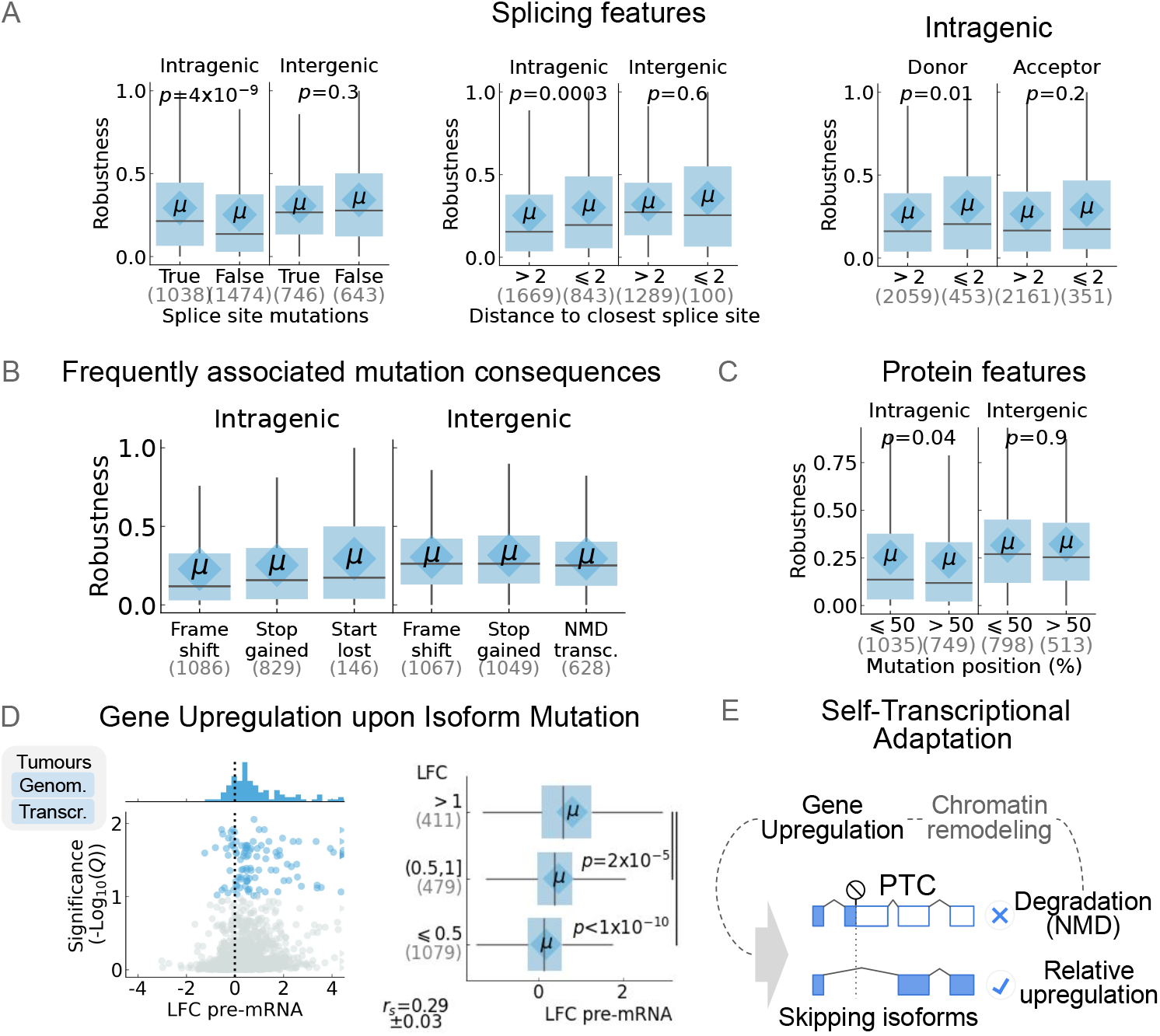
Regulatory mechanisms associated with mutational robustness. **A**: Association of mutation robustness scores with splicing-related features. Mutations are stratified based on whether they are located at the splice site (left), based on proximity to splice site (middle), or based on proximity to donor vs acceptor splice sites (right). In all box plots in this figure, *p*-values are obtained from two-sided Mann-Whitney U-tests. **B**: Top three most frequent mutation consequences (x-axis) among robust instances (robustness score>0) shown with corresponding robustness scores for intra and inter-genic levels (y-axis). **C**: Association of mutation position in protein products (first or second half of the protein, x-axis) with robustness scores for intra and inter-genic levels (y-axis). **D**: Transcriptional upregulation, measured as LFC of pre-mRNA, is shown as a volcano plot on left, and its association with LFC of the skipping isoforms is shown on the right. Q: adjusted *p*-value obtained from the differential expression analysis. *r*_*s*_: Spearman’s correlation. **E**: Schematic representation depicting the regulatory mechanism of self-TA.

While disruption of both donor and acceptor sites can theoretically result in exon skipping, as a key differentiating factor, donor site disruption in particular is more likely to lead to intronic readthrough and thus acquisition of premature termination codon (PTC) (**Fig S8**). This led us to hypothesize that, in addition to splice-switching mutations, acquisition of PTCs can also lead to a shift in isoform abundances post-transcriptionally, even when the underlying splicing ratios remain the same. Consistent with this hypothesis, overall, we observed that frameshift and stop-gain mutations (which generated PTCs) were the most frequent mutational events associated with intra-genic compensation (intra-genic robustness score>0) by alternative isoforms (**Fig 4B**). At protein level, mutations in the first half of protein length are associated with significantly higher intra-genic (but not inter-genic) robustness score compared to mutations in the second half of the protein (*p*=0.04, **Fig 4C**). Put together, these observations raised the possibility for the involvement of nonsense-mediated mRNA decay (NMD), triggered by PTCs ^45^, in the mechanism underlying intra-genic mutational robustness.

NMD has been proposed to lead to up-regulation of related genes via transcriptional adaptation, mediated by a mechanism that depends on the sequence similarity between the intermediate products of NMD and the sequence of adapting genes ^46–50^. Importantly, this adaptation can also happen at the perturbed gene itself, a mechanism referred to as self-transcriptional adaptation (self-TA) ^50^. We asked whether self-TA could be involved in intra-genic mutational robustness. We obtained gene-wise pre-mRNA expression profiles for the tumor datasets ^51^ and carried out differential analysis between the same groups of samples that were used to estimate the upregulation of the mutation-skipping isoforms (see **Methods**). Indeed, we found that somatic mutations are often associated with an increase in pre-mRNA abundance in affected samples (compared to cancer type- and expression profile-matched samples) (**Fig 4D**-left), including 178 events that showed a positive LFC (compared to 20 that showed a negative LFC, FDR<0.1; binomial P<1e-10). Importantly, we observed a highly significant correlation between pre-mRNA up-regulation and up-regulation of mutation-skipping isoforms (*r*_*s*_=0.29, *p<*1e-10), with genes that show the strongest mutation-skipping isoform up-regulation also showing the strongest pre-mRNA up-regulation (**Fig 4D**-right). Notably, this trend was found to be unique to genes with mutation-skipping isoforms; for the genes where none of the isoforms skipped the mutation, a non-significant association was observed (**Fig S9**).

Putting these findings together, a mechanism based on transcriptional adaptation ^46,52^ emerges where the isoforms carrying PTCs undergo NMD, which triggers upregulation of the transcription of perturbed gene. As the affected isoform(s) are degraded by NMD, this self-TA mechanism effectively leads to the relative upregulation of the mutation-skipping isoforms (**Fig 4E**).

### Implications of intra-genic mutational robustness

Next, we investigated how intra-genic robustness may affect the molecular functions of the corresponding genes. We specifically assessed protein-protein interactions (PPIs) since a gene’s functions are generally carried out via interaction with other proteins ^53^. We expected that PPI rewiring may be observed in response to up-regulation of mutation-skipping isoforms, in order for the perturbed gene to remain functional. Specifically, we tested isoform-level PPI rewiring using differential isoform usages (DIUs) of the perturbed gene and its interactors, restricting to PPIs within protein complexes since they tend to be cognate, typically exhibit a high experimental confidence, and tend to be sensitive to the stoichiometry of complex members (see **Supplementary Methods** for details). First, we compared the DIUs of the genes that are perturbed by a mutation, genes that interact with a perturbed gene, or neither. We expectedly observed that the ‘perturbed’ category exhibited higher DIU compared to ‘neither’ (*p*=1e-10, **Fig 5A**-left). Remarkably, ‘interactors’ too showed higher DIUs than the ‘neither’ category of genes. This pointed to the possibility of co-occurrence of DIU among interactors to accommodate the expression of mutation-skipping isoforms of the perturbed gene. Indeed, we found a significant correlation between the DIU of perturbed genes and DIUs of its interactors (*r*_*s*_=0.14±0.02, *p*=5e-10, **Fig 5A**-right), indicating co-ordinated isoform remodeling among interactors and suggesting restorative PPI rewiring at isoform-level.

**Figure 5:**
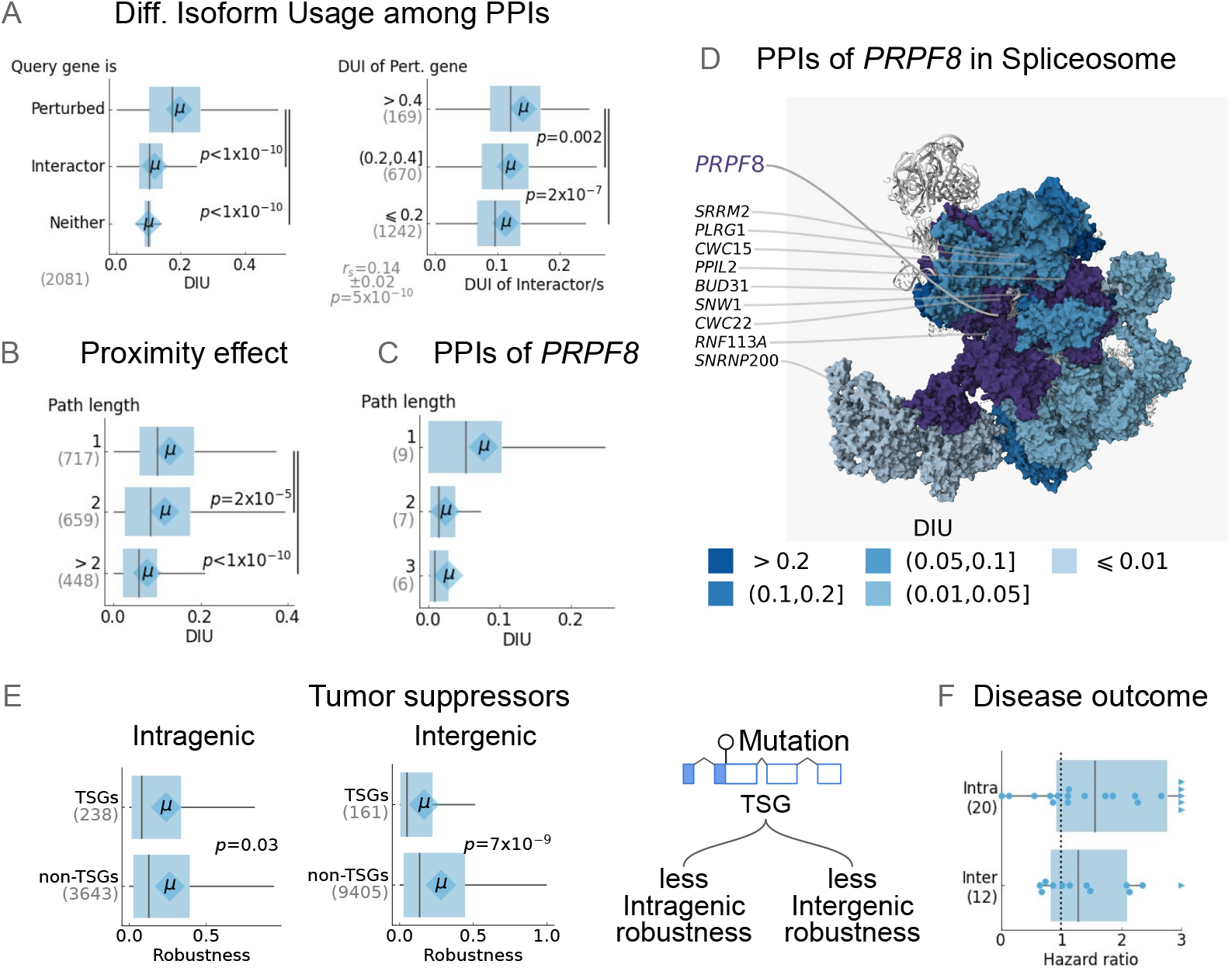
Implications of mutational robustness. **A**: Differential Isoform Usage (DIU) among PPIs belonging to protein complexes. Left: distribution of DIUs for the same genes is compared between contexts where they are ‘perturbed’, they are ‘interactors’ of perturbed genes, or ‘neither’ (control). Right: DIUs of perturbed genes are correlated with DIUs of corresponding interactors. *r*_*s*_: Spearman’s correlation coefficient. In all box plots in this figure, *p*-values are obtained from two-sided Mann-Whitney U-tests. **B**: The DIUs are compared between the interactors at different levels of proximity measured in terms of the shortest path lengths from the perturbed gene. **C**: Similar to panel B, but for the spliceosome subunit *PRPF8* as an example, which is perturbed in prostate adenocarcinoma cancer and exhibits intra-genic mutational robustness. The DIUs of its spliceosome interactors are compared between categories of shortest path lengths relative to *PRPF8*. **D**: DIUs of individual interactors of *PRPF8* are mapped onto the structure of spliceosome protein complex (PDB ID: 8i0s ^54,55^). The *PRPF8* and its proximal interactors are annotated. **E**: Mutational robustness scores among TSGs compared against non-TSG genes, for the intra-genic (left) and inter-genic level (middle). The resultant model is depicted on the right. **F**: Relationship between mutational robustness of non-TSGs and patient survival tested using Cox regression. Hazard ratios (x-axis)—i.e. the effect sizes (indicating risk) obtained from Cox regression—are shown for intra and inter-genic levels (y-axis). Each dot is a cancer type. Hazard ratios are corrected for age, gender, mutation burden, and race (see **Supplementary Methods**).

Furthermore, stratifying the interactors by their interaction path length from the perturbed gene, we found a clear trend of decreasing DIU with the increasing path length (*p*=9e-25, **Fig 5B**). An example is shown in **Fig 5C-D**, depicting intra-genic compensation of deleterious mutations in the *PRPF8* subunit of the spliceosome and the accompanying isoform remodeling of other spliceosome components. Specifically, in prostate adenocarcinoma, we observed up-regulation of a mutation-skipping isoform (ENST00000703540) in response to perturbation of four other isoforms by deleterious mutations. We observed a consistent trend of DIU elevation in its proximal interactors with dissipation onwards, up to the path length of three (**Fig 5C**), with the greatest DIU observed in direct PRPF8-interacting proteins (**Fig 5D**) involved in RNA anchoring (*BUD31* and *PPIL2*), coordinated helicase activities (*SNRNP200)*, spliceosome stabilization (*RNF113A*), and active site formation and splicing fidelity (*SRRM2, PLRG1, CWC15, SNW1* and *CWC22*).

We also investigated the relevance of mutational robustness to cancer development and outcomes. Cancer genomes most frequently harbor deleterious mutations in Tumor Suppressor Genes (TSGs) during early stages of cancer initiation and development; we asked whether such mutations are also compensated through inter- and intra-genic robustness pathways. Comparing intra-genic robustness scores between the TSGs and non-TSGs—in a cancer type-wise manner—we find that TSGs tend to have significantly lower robustness scores (*p*=0.03, **Fig 5E**-left). The same trend was also observed in the case of inter-genic mutational robustness (*p*=7e-9, **Fig 5E**-middle), indicating a general compromise of mutational robustness among TSGs. This finding suggests that beyond merely carrying the deleterious mutations, TSG functions tend to be inactivated in cancer cells by a lack of protective robustness. In other words, this finding suggests that, in addition to the TSG mutation itself, cancer cells must bypass inter- and intra-genic robustness mechanisms specifically for TSGs (**Fig 5E**-right). In contrast, robustness of non-TSG genes must be advantageous for cancer cells as it allows them to tolerate genomic instability. Consistent with this hypothesis, we found that in the majority of cancer types, high mutational robustness of non-TSGs, through intra- or inter-genic compensatory up-regulation, is associated with worse disease outcome (hazard ratio >1 for 16 out of 20 cancer types for which intra-genic robustness could be estimated, and 8 out of 12 cancer types for which inter-genic robustness could be estimated; **Fig 5F**). These observations suggest that while TSGs are preferably inactivated in cancer cells by compromised robustness, the functionality of non-TSGs tends to be protected by robustness, likely because non-TSG functions play an important role in cancer cell survival and disease progression.

## Discussion

Here, we present an expanded framework for mutational robustness in cancer in which functional compensation occurs recursively across nested molecular levels. Specifically, we show that alternatively spliced isoforms frequently bypass deleterious mutations and undergo compensatory up-regulation, thereby providing intra-genic mutational robustness that can further be complemented and augmented by inter-genic compensation through paralogs. Our analyses across tumors and cancer cell lines reveal that mutation-skipping isoforms are prevalent, exhibit compensatory expression dynamics associated with perturbation tolerance, and are accompanied by coordinated remodeling of interacting partners at the isoform level. Mechanistically, our findings support a model in which NMD of perturbed isoforms induces self-TA, resulting in relative up-regulation of mutation-skipping isoforms. Collectively, these observations expand the current view of “function-sharing” between isoforms and paralogs ^22–24^, and suggest that mutational robustness in cancer emerges through layered compensatory processes operating across multiple biological scales. More broadly, our findings suggest that mutational robustness in cancer is not solely a passive consequence of functional redundancy, but also involves active adaptive responses to perturbation. The prevalence of recursive mutational robustness further provides a plausible explanation for the remarkable tolerance of cancer cells to pervasive genomic perturbation despite largely asexual evolution and limited opportunities for purifying selection ^1–4^.

We also showed that this robustness landscape was strongly associated with expression level, consistent with the concept of need-based robustness. Genes exhibiting strong intra-genic or inter-genic robustness tended to be highly expressed, suggesting that compensation preferentially occurs where preservation of gene function is most required. Indeed, highly expressed genes are often dosage-sensitive and evolutionarily constrained ^42^, creating selective pressure to preserve their functional output. A plausible mechanistic explanation is provided by self-TA, given that highly expressed genes are expected to produce larger quantities of NMD degradation intermediates capable of triggering transcriptional adaptation ^46,48,49^. Additional mechanisms may further reinforce this pattern, including permissive chromatin environments that may facilitate rapid compensatory transcriptional responses, and broader expression scaling mechanisms (such as autoregulatory circuitry) that maintain stable biomolecular concentrations ^28,29^. Need-based robustness may, therefore, emerge through convergence between molecular sensing capacity and selective pressure to preserve essential cellular functions.

Our analyses further revealed that genes exhibiting strong intra-genic robustness also tended to exhibit stronger inter-genic compensation through paralogs, giving rise to what we refer to as recursive mutational robustness. One explanation is that both mechanisms share a common upstream mechanism, i.e., perturbation-induced transcriptional adaptation; thus, genes that strongly activate compensatory responses at the isoform level may also be more likely to induce compensatory up-regulation of paralogs. However, recursive robustness may additionally reflect deeper organizational principles of cellular systems, with selective pressures on genes with dosage-sensitive functions ^56^ leading to accumulation of multiple overlapping compensatory architectures through evolution.

We note that, although compensatory up-regulation was the dominant response to perturbation, we also observed rare instances in tumour data in which mutation-skipping isoforms were down-regulated instead (Fig 2B), with more such examples in CRISPR perturbation data (Fig S6A). As we showed in Fig 3E, these events were enriched among lowly expressed genes in the CRISPR perturbation data, consistent with weaker activation of need-based compensatory responses. In some cases, larger-than-expected CRISPR genomic alterations may additionally disrupt all isoforms simultaneously, thereby preventing selective compensation. Another alternative explanation is the existence of isoform co-dependency as a potential mechanism opposing compensation-based robustness, similar to previously reported gene level co-dependency ^7^. Consistent with this potential mechanism, we saw that genes exhibiting down-regulated mutation-skipping isoforms showed stronger coexpression among isoforms (Spearman correlation of –0.65 between isoform coexpression and LFC of mutation-skipping isoform, p=0.008; **Fig S6C**), suggesting that some isoforms function cooperatively rather than redundantly. Such dependencies may be particularly relevant among RNA-binding proteins, where autoregulatory feedback loops and coordinated expression programs are common ^57^.

Interestingly, while mutational robustness was widespread among non-TSGs, it appeared substantially compromised among TSGs. This observation suggests that successful tumor suppressor inactivation may require not only acquisition of deleterious mutations, but also evasion of compensatory robustness mechanisms. One possibility is that robustness constrains the subset of mutations capable of functionally inactivating TSGs. For example, mutations targeting constitutive exons of TSGs may be more effective because they cannot readily be bypassed by alternative splicing and, thus, are more abundantly present in cancer cells, whereas mutations affecting dispensable alternative exons may remain buffered through intra-genic compensation. Similarly, among multiple genes capable of suppressing a given oncogenic pathway, genes with intrinsically weaker compensatory architectures may be preferentially selected during cancer evolution because they are more readily inactivated. More broadly, these observations raise the possibility that mutational robustness itself shapes the accessible evolutionary trajectories of tumor suppressor inactivation in cancer.

Finally, an important implication of intra-genic mutational robustness is that it may create highly selective therapeutic vulnerabilities. In cancer cells harboring deleterious mutations, mutation-skipping isoforms can become functionally required to maintain gene activity, effectively shifting dependence from the perturbed isoform toward compensatory isoforms. This creates a potential isoform-level synthetic lethality framework in which selective inhibition of mutation-skipping isoforms could preferentially impair cells carrying the corresponding mutation, whereas cells lacking the mutation would be expected to better tolerate inhibition of the skipping isoforms as the intact original isoform would remain available to preserve gene function. Such dependencies may be particularly amenable to therapeutic targeting using isoform-selective approaches, including antisense oligonucleotides directed against specific exon-exon junctions ^58^. More broadly, these observations suggest that mutational robustness is not only a mechanism of tolerance, but may also expose selective liabilities that emerge specifically from compensatory adaptation.

## Methods

### Genomic annotations

For uniformity, all the isoform-level analysis done in study uses the same version of the genome annotations (GENCODE V46, Ensembl release 112). See **Supplementary Methods** for details.

### Integration of the genomics and transcriptomics data

Somatic mutations and RNA-seq data for tumors were obtained from TCGA (https://www.cancer.gov/tcga) whereas cell line data were obtained from CCLE ^32^. Briefly, both datasets were mapped to the same genome annotation version. Isoform-level expression was quantified using Salmon ^59^, and deleterious mutations were detected using Ensembl VEP version 112 ^60^. We developed a classification system (described in **Supplementary Methods**) to assign the mutation consequence to the corresponding isoforms and mark them as deleterious. In the corresponding gene-wise dataset, the isoform-wise mutation status and expression were aggregated by gene IDs. The resultant datasets encompassed a total of 7307 tumors and 340 cell lines. More details are provided in **Supplementary Methods**.

### Percentage of perturbation skipping isoforms per gene

To quantify the proportion of skipping isoforms per gene across given samples, we first retained only the genes with multiple isoforms (≥ 2). We mapped them to the given isoform-wise binarized modality (*m*) of interest: mutation status (any mutation), carrying deleterious mutation, or expressed (TPM≥ 1). For each sample, we first calculated the percentage (*p*_*affect*_) of affected isoforms (total = *I*) per gene. Then, these sample-wise values were averaged across samples with at least one affected isoform, preventing a zero-inflation bias. Subtracting this value from 100 provided a single score for a given gene indicating the averaged percentage of skipping isoforms per gene (*p*_*skip*_).

### Mutational robustness scores

#### Input datasets

We divided the tumor data by the cancer type and cell line data by lineages. We only retained such sample subsets with a sufficient statistical representation (≥3 samples). The datasets stored in multimodal data container format (.h5mu) were provided to a workflow we developed, called “intom” (https://github.com/csglab/intom). The workflow calculates robustness scores in each sample subset (as described below). The fusion genes reported in tumor data (TCGA) or cell line databases (DepMap) were excluded from the analysis.

#### Formulation

As shown in **Fig2 A**, isoforms are intra-genic entities grouped by gene, while paralogous genes are inter-genic entities grouped by paralog family. Note that the mutational robustness is an interaction between perturbed and perturbation-skipping entities. The samples in which the same set of entities were perturbed are considered the ‘test’ samples. They were compared against ‘reference’ samples that (a) were from the same sample group, (b) did not contain any mutations in any entities, and (c) had a close similarity in terms of expression profiles, based on Pearson’s correlation coefficient (*r*_*p*_) ≥0.9 across all genes. We only retained comparisons with a sufficient number of reference samples (at least 3). These reference samples served as the control group in downstream differential analysis. Note that, in the case of inter-genic interactions, even if robustness could be traced down to an individual dominant isoform ^61^ of the compensating gene, following previous studies ^62^ we still consider this an inter-genic interaction by the virtue of the involvement of multiple genes.

#### Backup upregulation

We measured the differential expression of the mutation-skipping entities between ‘test’ and ‘reference’ samples, using empirical Bayes variance shrinkage through limma ^39^—for each comparison, the limma model was fit to all genes/transcripts, to enable proper estimation of model parameters, but only the relevant statistics for the perturbed on perturbation-skipping entities were extracted. For the skipping entities, LFC represented the extent of up/down regulation and the adjusted *p*-values indicated the corresponding statistical significance level. We filtered the differentially expressed instances to retain the ones with non-spurious expression levels (mean log2-TPM ≥0.1 across samples included in the comparison).

#### Compensation

Focussing on upregulation of skipping instances, we assessed whether the extent of upregulation would be sufficient for carrying out functional compensation. For this, we calculate the compensation score as the ratio between the total expression of the skipping entities, averaged across ‘test’ samples, and the total expression of all the entities, averaged across the ‘reference’ samples. For example, if an isoform that skips a mutation has a mean expression of 1 TPM in samples carrying that mutation, and the corresponding gene has a mean expression of 2 TPM in reference samples (without the mutation), the compensation score is 1/2=0.5.

#### Robustness scores

Finally, the robustness scores scores were calculated by multiplying the extent of upregulation and compensation, both thresholded between 0 and 1 to mitigate the influence of extreme values. This multiplicative approach ensures that a high robustness score requires both strong upregulation and a substantial amount of compensation. For example, a robustness score close to 1 requires both LFC close to or greater than 1 and compensation close to or greater than 1 (see **Fig S3A**). Note that since the upregulation LFCs are calibrated against the noise level—owing to the empirical Bayes variance shrinkage through limma ^39^—use of the scores was preferred over thresholding by statistical significance of LFCs (Q-values). This prevented inducing possible truncation and data availability biases.

#### Gene set enrichment

We obtained Gene Ontology (GO) terms for Biological Process, Molecular Function and Cellular Component aspects, corresponding to the genome annotation used in the study, through BioMart. The extremely small and extremely large gene sets (<20 or > 3000 genes) were not considered. The gene sets were mapped to the genes with moderate-to-high robustness score (> 0.25). To account for potential confounding factors, we incorporated gene-level covariates: gene length, average mRNA expression level (from tumor dataset), and the expression entropy (from tumor dataset) in a logistic regression model evaluating the overrepresentation (odds ratio > 1) of the genes with high robustness among the gene sets.

### Transcriptomic profiling with CRISPR-based perturbation

CRISPR RNA-seq datasets and sgRNA metadata for K562 and HepG2 cell lines were retrieved from ENCODE ^43^. Targeted isoforms were strictly filtered to classify isoforms as perturbed if the sgRNA aligned within 50 bp of an exon boundary. Additional details can be found in **Supplementary Methods**.

## Data and code availability

The code for the analyses shown in the manuscript is available at https://github.com/csglab/robustness_ms, and the associated data for reproducible analysis can be found at Zenodo (DOI: 10.5281/zenodo.20382591). The intom workflow is available at https://github.com/csglab/intom.

## Acknowledgements

We thank the CSG lab for discussions and for providing feedback on our manuscript. This work was supported by grants from the Canadian Institutes of Health Research (CIHR) [PJT-173317] to HSN, and resources provided by Calcul Québec (calculquebec.ca) and the Digital Research Alliance of Canada (alliancecan.ca) to HSN. HSN holds a CIHR Canada Research Chair. HG is an Arc Institute Core Investigator.

## Author contributions

RD: Conceptualization, Methodology, Software, Validation, Formal analysis, Investigation, Data Curation, Writing - Original Draft, Writing - Review & Editing, Visualization; AHC: Methodology, Writing - Review & Editing; AM: Methodology, Writing - Review & Editing; AMN: Methodology, Writing - Review & Editing; JW: Methodology, Writing - Review & Editing; BC: Methodology, Validation, Writing - Review & Editing; HG: Methodology, Validation, Investigation, Resources, Writing - Review & Editing, Supervision; HN: Conceptualization, Methodology, Investigation, Resources, Writing - Original Draft, Writing - Review & Editing, Supervision, Project administration, Funding acquisition.

