## Supplementary Information for "Recursive mutational robustness in cancer through intra- and inter-genic compensation"

### Supplementary figures

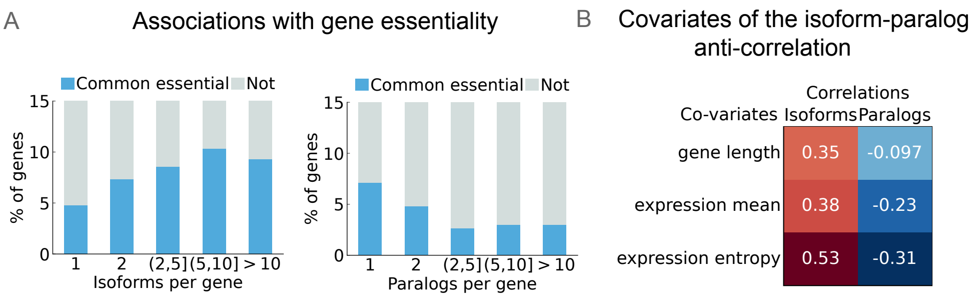


**Figure S1: Properties of genes with isoforms and paralogs.**
**A**: The percentage of common essential genes (y-axes) among genes with different numbers of isoforms (left) and paralogs (right).
**B**: Covariates of the isoform-paralog anticorrelation. On the left, the Spearman correlation coefficients ($r_s$) between the number of isoforms and paralogs with covariates: expression entropy (based on the isoform expression probability per gene), gene expression (mean across cancer types of tumors) and gene length are shown.

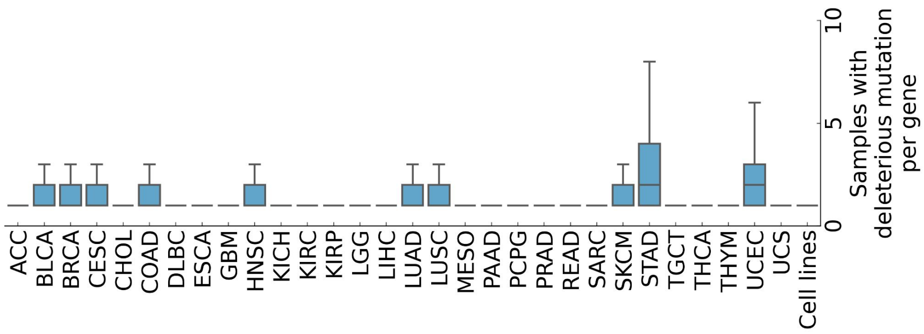


**Figure S2:** Distributions of samples with deleterious mutations per gene across the tumour types and cell lines analyzed in the study.

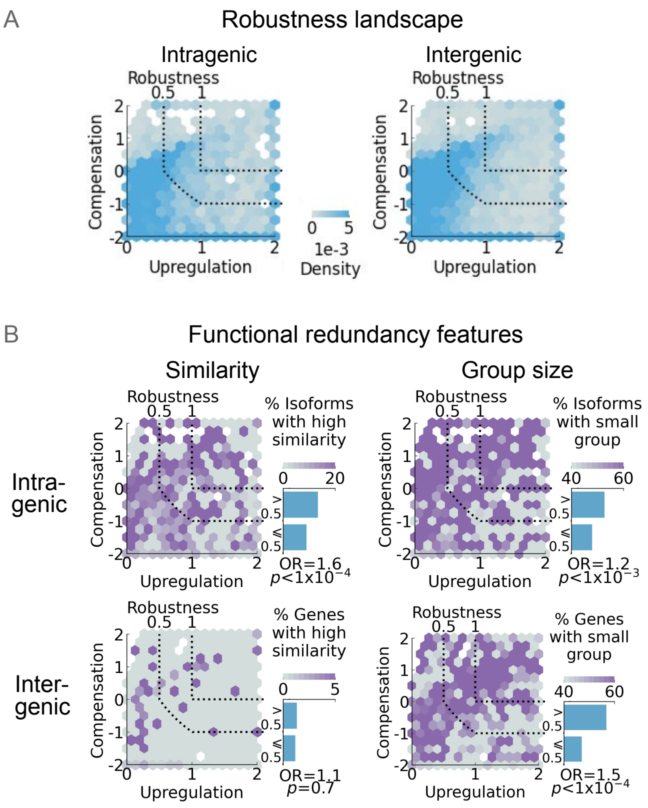


**Figure S3: Robustness landscape visualization and the functional redundancy features mapped onto it.**
**A**: Robustness landscape visualization showing the density of intragenic and intergenic mutational robustness instances lined by upregulation and compensation. For visual clarity, compensation is shown on log-scale.
**B**: The intragenic (top row) and intergenic (bottom row) robustness landscapes mapped by the percentage of entities (shown with the color gradient) with high similarity (protein sequence similarity max. per group >90%, left), and a small group (group size<10, on right). The bar plots show the associations with high (>0.5) and low robustness scores (≤0.5). OR: Odds ratio, *p*: *p*-value associated with Fisher's exact test.

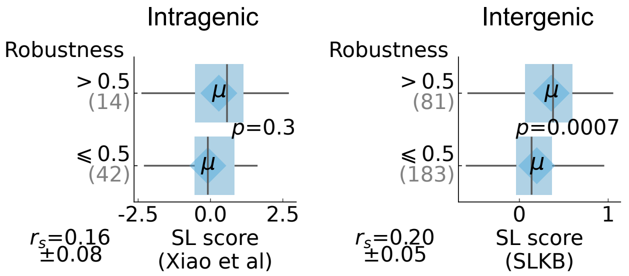


**Figure S4: Associations of the mutational robustness scores (y-axis) with tolerance of CRISPR-based perturbations (x-axis).**
We used synthetic lethality (SL) scores estimated using exon-deletion screen data from Xiao et al. for intragenic level, and paired gene deletion data from the SLKB dataset[1] for intergenic level analysis (See **Supplementary Methods** for details). In both cases, higher scores represent higher tolerance of perturbation. The associations were assessed using correlation as well as a binned approach comparing the measure of tolerance (x-axis) between the two robustness score bins (y-axis). Note that since SL data originated from cell lines, it was compared against the robustness scores of the cell lines datasets. *p*: *p*-value obtained from two-sided Mann-Whitney U-test; $r_s$: mean Spearman's correlation across 5 resamplings, with min/max confidence interval. See Supplementary Methods for detailed Methods.

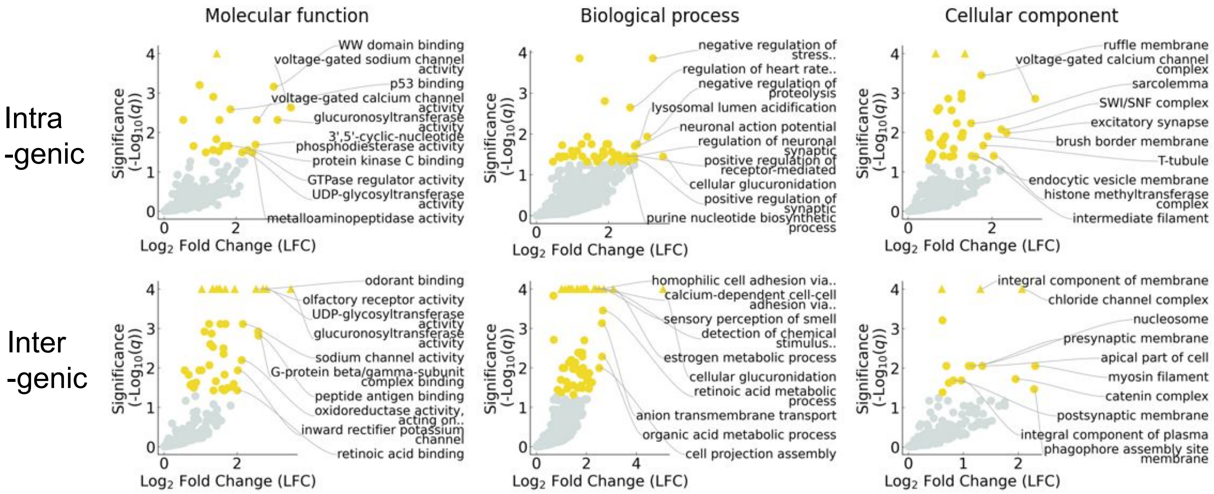


**Figure S5:** Enrichments of Molecular functions (left), Biological process (middle) and Cellular Component (right) gene-sets among genes exhibiting mutational robustness (robustness score>0) at intra- (top) and intergenic (bottom) levels. The top ten gene sets with the greatest enrichment and significance are annotated. LFC: $Log_2$ Fold Change; q: adjusted p-value obtained from logistic regression.

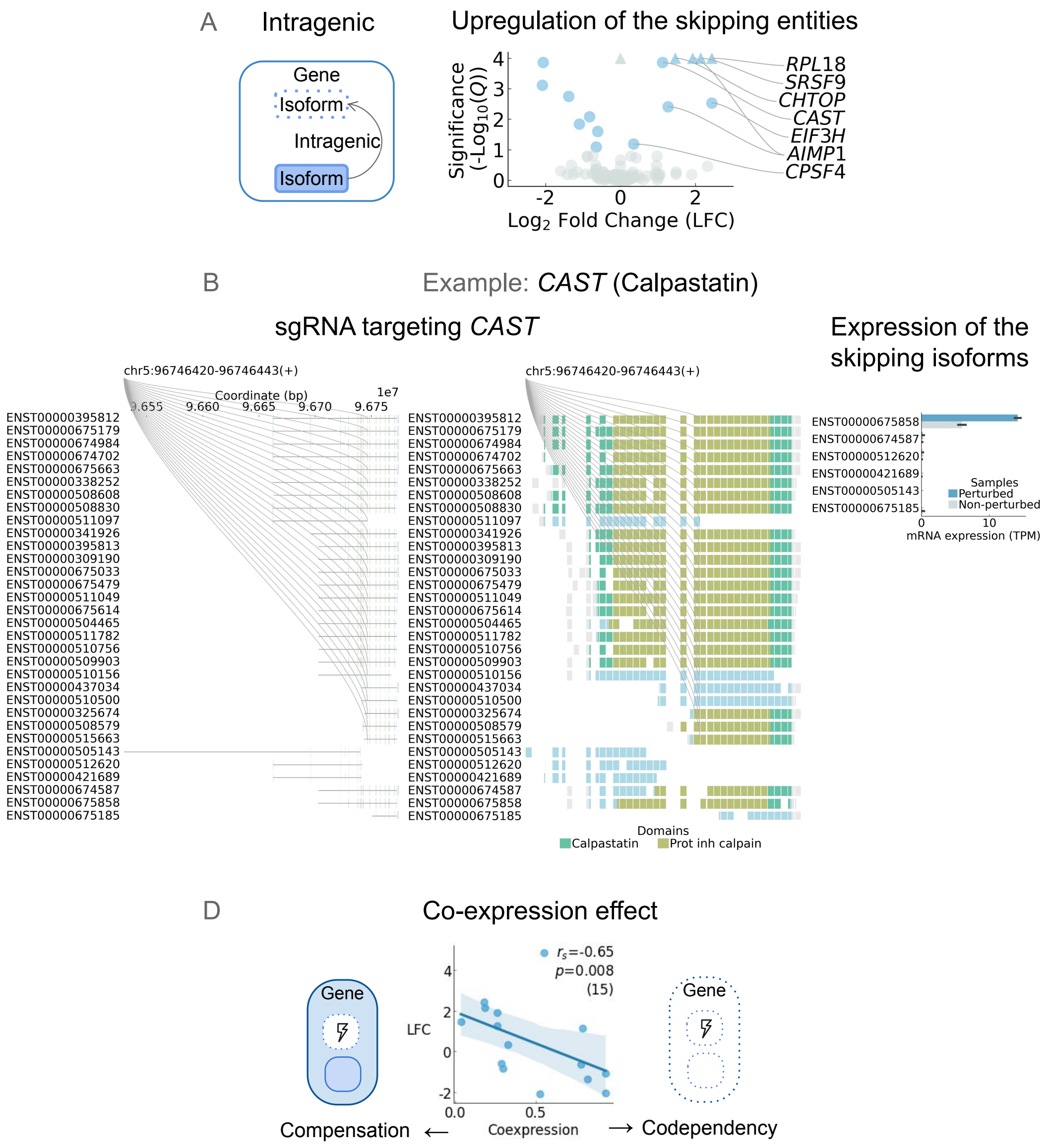


**Figure S6: Intra-genic mutational robustness in response CRISPR perturbation.**

**A**: Left: schematic representation of the intragenic robustness interaction. Right: differential expression of perturbation-skipping isoforms.

**B**: Intra-genic robustness in *CAST* (Calpastatin) gene. Left: sgRNA positions are shown along the genomic coordinates and, for visual clarity, on the exon blocks. Right: differential expression of the skipping isoforms between the samples where the CRISPR perturbation of the gene was carried out versus the samples without the CRISPR perturbation.

**C**: Co-expression between isoforms (x-axis) correlated against the LFC of the statistically significant instances of intragenic mutational robustness (y-axis). $r_s$: Spearman's correlation coefficient, $p$: associated $p$-value. Trend lines are fitted with linear regression, and their 95% confidence interval is shown.

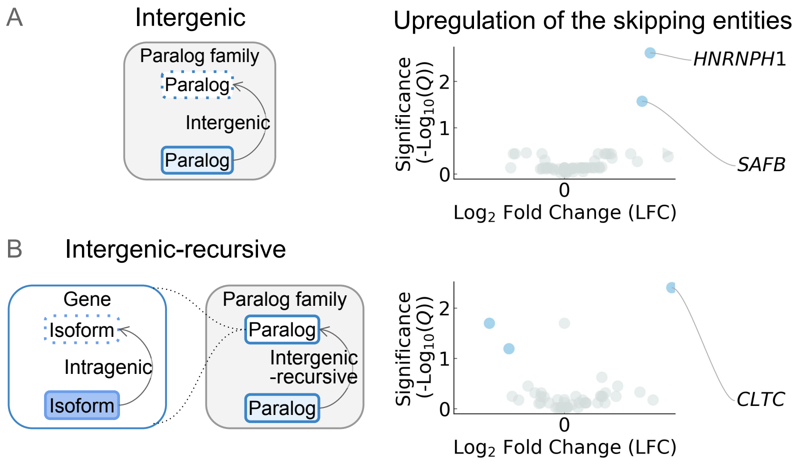


**Figure S7: Intergenic and recursive mutational robustness for the CRISPR perturbation.**
**A**: Similar to Fig S6A, except that the 'integenic' setting is shown.
**B**: Similar to Fig S6A, but for the 'recursive' setting.

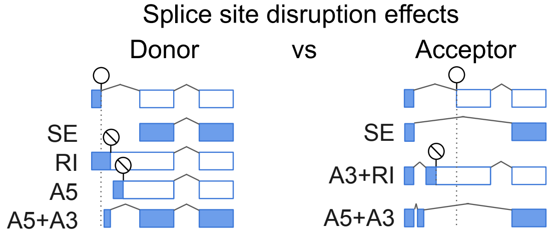


**Figure S8:** Schematic representation of likely outcomes from donor versus acceptor splice site disruptions, including skipping the exon (SE), usage of alternative 5' or 3' site (A5 and A3 resp.), and retained intron (RI).

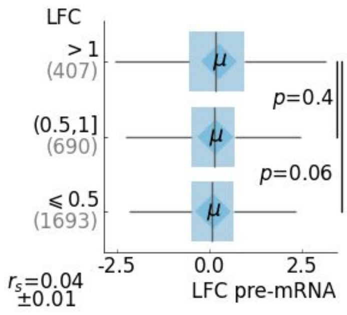


**Figure S9: Differential pre-mRNA expression for genes without perturbation-skipping isoforms.**
Similar to **Fig 4D**-right.

### Supplementary methods

#### Genome annotations

The gene annotations for protein-coding genes were obtained from BioMart (GENCODE V46, Ensembl release 112) using the pyBiomart interface 2. These annotations included 20065 protein-coding genes, with 89581 protein-coding isoforms and 458982 exon annotations. The protein domain annotations corresponding to the same version were obtained from the BioMart.

#### Isoforms

Sequence similarity: For each gene, isoform-wise sequence similarities between the protein-sequences was obtained using Needle [3] with the following parameters: EBLOSUM62 -gapopen 10.0 -gapextend 0.5 -endopen 10.0 -endextend 0.5 -aformat3. The sequence similarity cutoff of minimum 75% was used to categorize isoforms with similar sequences.

Alternative splicing events: The alternative splicing events corresponding to the isoforms were assigned using SUPPA2 [4].

#### Genes

Paralogous genes: The pairs of paralogous genes were obtained from BioMart for the genome annotation version used in the study, using the pyBiomart interface. The sequence similarities between the protein products of the paralogs were calculated in a similar manner as isoforms. The paralog pairs with a minimum sequence similarity of 50% were retained, to ensure sufficient number of pairs for subsequent analyses.

Paralog families: The paralog families were assigned to the paralogs by identifying the connected components in a graph generated by using paralogs as nodes and pairs as edges.

#### Gene ontology terms

GO-terms were obtained from BioMart for the genome annotation version used in the study, using the pyBiomart interface.

#### Cancer genes

Annotations of the Tumor-Suppressor Genes (TSG) were obtained from the COSMIC Cancer Gene Census (v98, for GRCh38)[5].

#### Integration of the genomics and transcriptomics data

Data retrieval and pre-processing: For tumours, Whole Genome Sequencing (WGS)-based somatic mutation data in VCF format (wgs.GATK4_MuTect2_Pair.somatic_annotation.vcf) and corresponding RNA-seq data in BAM format were obtained from TCGA (https://www.cancer.gov/tcga), through GDC Data Portal (https://portal.gdc.cancer.gov, downloaded on 2025-09-28). For cancer cell lines, mutations were obtained in the Mutation Annotation Format (MAF) from the DepMap portal (DepMap, release: 23Q4) [6]. Data from Whole-Genome Sequencing (WGS) was used. The co-ordinates were mapped to the version of the gene annotations used in the study using bedtools's [7] intersect functionality. Cell line RNA-seq data were obtained from the Cancer Cell Line Encyclopedia (CCLE) [8]. Specifically, we used the fastq files obtained from Sequence Read Archive, SRA (BioProject ID: PRJNA523380).

#### Data processing

Variant effect prediction: The variant effects were predicted using Ensembl VEP 112 (McLaren et al. 2016) using default settings. Ensembl VEP was installed using Docker image (https://hub.docker.com/r/ensemblorg/ensembl-vep).

Classification of the deleterious mutations: Using the predicted variant effects, mutations annotated with following the Sequence Ontology (SO) terms were classified as deleterious: stop_gained, frameshift_variant, transcript_ablation, feature_truncation, splice_acceptor_variant, splice_donor_variant, protein_altering_variant, and start_lost. Mutations annotated with the following SO terms were classified as deleterious only if the associated impact is HIGH: NMD_transcript_variant, coding_sequence_variant, splice_region_variant, splice_donor_region_variant, and splice_donor_5th_base_variant.

mRNA expression: The mRNA expression in TPM units was quantified using salmon [9]. Nextflow rnaseq workflow [10] version 3.15.1 was used to carry out this analysis, using default settings for the pre-processing of the raw data.

Gene-wise data: Isoform-wise deleterious mutation status and expression was mapped to gene IDs using the same genome annotation version. Deleterious mutation status (boolean) was aggregated to gene-wise percentages of isoforms carrying the mutation. Isoform-wise expression was summed per gene to get the gene-wise expression.

**Fitness effects of CRISPR-based perturbation**

Gene deletion screen data: The gene essentialities estimated from CRISPR-based gene inactivation screens were obtained from the DepMap portal (DepMap, release: 23Q4)[6]. The gene effect scores were used as a measure of gene essentialities. Common essential genes reported by the same source, i.e., DepMap, were used.

Exon deletion screen data: The exon essentiality data measured from CRISPR-based screens was obtained from Xiao et al. [11]. We re-annotated the genome-wide exon deletion screen data to map with the transcript model version used in this study.

Fitness effects: To evaluate the fitness effects of exon deletions, we processed exon-level functional screen data. We isolated in-frame deletion events within the exon deletion library. For each exon, we calculated the mean fitness effect (exon beta) across the cell lines. Concurrently, we aggregated gene-level fitness data to identify genes essential in at least one sample. To classify the functional impact of individual exon deletions, we restricted our thresholding analysis strictly to exons located within these high essentiality genes. We fitted a two-cluster one-dimensional Gaussian Mixture Model (GMM, shown in Fig 1E left) to the mean exon beta scores to obtain a statistical cutoff to classify exons into fitness decrease and neutral categories.

**Comparison of the homozygous deletion frequencies between exon types**

First, to identify homozygous deletions, we processed structural variant data from TCGA paired tumor-normal whole-genome sequencing (downloaded on 2025-09-22). We retained precise variants (IMPRECISE ≠ True) with high-confidence homozygous deletion support (TUMOR: homozygous score ≥ 0.75 and TUMOR: alt count ≥ 3). Variant coordinates were converted to a 1-based system and intersected with autosomal exon coordinates. We calculated the proportion of each exon overlapped by the deletion segment (segment overlap %). To remove potential artifacts, we excluded deletion loci overlapping with known genomic confounders, specifically fragile sites [12] and sequence gaps (https://hgdownload.soe.ucsc.edu/goldenpath/hg38). Finally, we aggregated the data to calculate exon-wise HD frequencies across samples and quantified the total number of exons spanned by each deletion segment per transcript.

Then to evaluate whether HD frequencies differ between exon types, we first mapped exon-level HD frequencies to exon types (constitutive or alternative). We isolated single-exon deletion events by filtering out multi-exon hits (>1 exon) and aggregated the sample-level HD frequency for each exon by taking mean. We restricted the analysis to multi-exon genes containing both exon types and strictly to exons with any occurrence of HD (i.e., frequency>0). We then calculated the mean mutation frequency by exon type within each gene and to allow stratification by gene type (paralog or singleton), mapped the gene-level frequencies to paralog annotations.

**Comparison of robustness scores**

Intragenic deleterious mutation frequencies: Using the isoform-level classification of the deleterious mutations for the TCGA whole-genome sequencing data (see above), we first calculated the frequency across all samples (samples with mutation/all samples). In the subsequent analysis, to mitigate the effect of extreme overrepresentation of the low frequency instances, we only retained the frequencies greater than 0.1% (> 0.001). These frequencies were mapped to the estimated robustness scores by perturbed isoforms and multiple robustness scores per isoform were aggregated by taking the maximum value.

Intergenic deleterious mutation frequencies: Similarly, using the gene-wise classification of the deleterious mutations for the TCGA whole-genome sequencing data (see above), gene-wise deleterious mutation frequencies were calculated. These values were mapped to the estimated robustness scores by perturbed genes and multiple robustness scores per gene were aggregated by taking the maximum value.

Intragenic SL score: To compute the intragenic SL score, — analogous to genetic interaction scores — we used exon deletion fitness effects processed from Xiao et al. study [11]. We first calculated isoform-wise fitness effects as the mean fitness effect (exon beta), across the perturbed exons and samples (two cell lines). For the corresponding genes, we calculated additive fitness effect as the sum of the individual isoform-level fitness effects. The gene-level mean fitness effect, — obtained from the same study, averaged across the same samples — served as the fitness effect upon the combined deletion of the gene's isoforms. Only the genes for which all isoforms possessed fitness effect scores were retained. Finally, SL scores were calculated by subtracting the (expected) additive fitness effect from the (observed) combined fitness effect, and multiplying by -1. These values were mapped to the estimated robustness scores in a gene-wise manner and multiple robustness scores per gene were aggregated by taking the maximum value.

Intergenic SL score: We obtained the SL scores across 10 CRISPR double-gene knockout screens, from the Synthetic Lethality Knowledge Base (SLKB, downloaded on Oct 14, 2025) [1]. This data covered 3417 paralog pairs and 21 cell lines. We calculated the mean SL scores and multiplied them by -1 for the paralog pair screened in multiple cell lines and screens. These values were mapped to the estimated robustness scores by gene pairs (perturbed and skipping entity) and multiple robustness scores per pair were aggregated by taking the maximum value.

**Transcriptomic profiling with CRISPR-based perturbation**

Publicly available CRISPR RNA-seq experiments data for K562 and HepG2 cell lines were retrieved from the ENCODE project portal [13](https://www.encodeproject.org, date: 2025-08-21). The dataset contained on average, 2 replicates per individual perturbation. The sgRNA sequences from the metadata were mapped to GRCh38 using Bowtie2 [14] — through alignment utilities of the beditor workflow [15], — and filtered for high-quality of alignment to single genes. Overlaps with Ensembl gene annotations used in the study were computed using bedtools intersect [7].

The identification of the targeted isoforms allowed for the identification of the isoforms which may skip the perturbation. This alignment-based "skipping" however may be compromised if the induced indel disrupts the immediate splice site consensus sequences, which can destabilize the splicing of the adjacent exons. We addressed this caveat by calculating the distances of the sgRNA sites from the nearest exon boundaries and marked the isoforms that were at the distance of <50 bp as perturbed. This included the isoforms with an intronic target site but at the proximity of the exon boundaries. This still leaves the possibility of large deletions extending into adjacent exons, but they tend to be rather rare compared to localized disruptions (max. 50bp on average) [16]. Another potential caveat is the disruption of intronic regulatory elements, prediction of which remains challenging[17]. However, considering that the splice site proximal regions — which we accounted for — tend to be more dense with SREs than the deep intronic spaces [18], the expression of the skipping isoforms is expected to be largely unharmed by the perturbation.

The filtered data contained candidate genes (counts shown in Fig 3D), for applying the robustness estimation method (`intom`).

The RNAseq data was used to obtain isoform and gene level expression quantification, using the same approach as for the tumor and cell line datasets. Expression and perturbation matrices were integrated at the isoform and gene level using MuData format [19].

#### Pre-mRNA expression

To quantify intronic read counts, we mapped the paired-end alignment data (BAM format) to reference annotations (GFF format) using featureCounts [20], through the CRIES workflow (https://github.com/csglab/CRIES) [21]. We restricted the feature summarization to intronic regions (feature type = intron) and obtained the mapped read counts at the gene level.

#### Differential Isoform Usage among PPIs

Differential Isoform Usage (DIU): To calculate DIUs, we first mapped the isoform-level expression profiles to the isoforms and genes and sample groups, for which the robustness score was scored. We aggregated the expression across samples in a given sample group by calculating the mean for each isoform. Missing or zero expression values were replaced with a pseudo-count to prevent downstream zero-probability vectors. We then calculated relative isoform usage profiles by normalizing the isoform means to sum to one for each gene within the respective sample groups. We quantified the per-gene isoform usage divergence by calculating the Jensen-Shannon distance between the paired sample groups. Finally, we filtered the dataset to exclude genes yielding undefined distance distributions and to ensure baseline transcriptional activity, we only retained the strictly expressed genes (TPM >= 1 in both sample groups).

Protein-protein interactions: The protein complexes and the PPIs within them were obtained from Complex portal [22] (downloaded on 22 Nov 2025). We excluded non-protein entities and self-interactions and retained only the interactors with valid UniProtKB identifiers and mapped them to Ensembl gene IDs. The shortest path lengths of the interactors from a given (perturbed) gene was measured using networkx Python library [23].

#### Survival analysis

The clinical and survival metadata for TCGA dataset were obtained from the Xena platform [24]. To the metadata of each cancer type, corresponding robustness scores (of intergenic or intragenic mutational robustness) were mapped and then binarised by sample IDs if any instance of robustness were found in the sample among non-TSG genes. Only primary tumors were considered. For fitting Cox's proportional hazard model, binarized robustness was used as the primary variable, age, gender, mutation burden, and race as covariates. If covariates had uninformative constant values (no variation) in a given regression model, they were removed. Additionally, regressions were performed only if the robustness variable was non-constant, sample size was >=20 and >=5 events were observed. The Submitter IDs were used as unique identifiers for clustering covariances. The models were fitted using lifelines Python package [25]. From the fitted models, the proportional-hazard ratios and corresponding *p*-values were obtained. To evaluate the association of the robustness scores with prognostic impact, among the fitted models we retained only the valid models satisfying the proportional hazards assumption.

#### Data analysis and visualizations

Table operations were carried out using the numpy [26] and pandas python package [27]. Statistical analyses were carried out using SciPy [28], and scikit-learn [29]. The network related analyses were carried out using networkx [23]. The data visualization was done using matplotlib [30], seaborn [31]. The Python package roux (https://github.com/rraadd88/roux) was used for both data analysis and visualization. The versions of these and other tools used in the analysis are provided with the source code. The isoforms were visualized using a Python package chrov [32].
